# Influenza A viruses use multivalent sialic acid clusters for cell binding and receptor activation

**DOI:** 10.1101/264713

**Authors:** Christian Sieben, Erdinc Sezgin, Christian Eggeling, Suliana Manley

## Abstract

Influenza A virus (IAV) binds its host cell using the major viral surface protein hemagglutinin (HA). HA recognizes sialic acid, a plasma membrane glycan that functions as the specific primary attachment factor (AF). Since sialic acid alone cannot fulfill a signaling function, the virus needs to activate downstream factors to trigger endocytic uptake. Recently, the epidermal growth factor receptor (EGFR), a member of the receptor-tyrosine kinase family, was shown to be activated by and transmit IAV entry signals. However, how IAV engages and activates EGFR remains largely unclear.

We used multicolor super-resolution microscopy to study the lateral organization of both IAV attachment factors and its functional receptor at the scale of the IAV particle. Intriguingly, quantitative cluster analysis revealed that AF and EGFR are organized in partially overlapping submicrometer clusters in the apical plasma membrane of A549 cells. Within AF domains, which are distinct from microvilli, the local AF concentration, a parameter that directly influences virus-cell binding, reaches on average 10-fold the background concentration and tends to increase towards the cluster center, thereby representing a multivalent virus-binding platform. Using our experimentally measured cluster characteristics, we simulated virus diffusion on a membrane, revealing that the distinct mobility pattern of IAVs is dominated by the local AF concentration, consistent with live cell single-virus tracking data. In contrast to AF, EGFR resides in clusters of rather low molecular density. Virus binding activates EGFR, but interestingly, this process occurs without a major lateral EGFR redistribution, instead relying on activation of pre-formed clusters, which we show are long-lived.

Taken together, our results provide a quantitative understanding of the initial steps of influenza virus infection. Co-clustering of AF and EGFR permit a cooperative effect of binding and signaling at specific platforms, and thus we relate their spatial organization to their functional role during virus-cell binding and receptor activation.

**Author Summary:** The plasma membrane is the major interface between a cell and its environment. It is a complex and dynamic organelle that needs to protect as a barrier but also process subtle signals into and out of the cell. For IAV, an enveloped virus, it represents a major obstacle that it needs to overcome during infection as well as the site for the assembly of progeny virus particles. However, the organisation of the plasma membrane in particular the sites of virus interaction at the scale of an infecting particle (length scales < 100 nm) remains largely unknown.

Sialic acids serve as IAV attachment factors but are not able to transmit signals across the plasma membrane. Receptor tyrosine kinases were identified to be activated upon virus binding and serve as functional receptor. How IAV engages and activates its functional receptors still remains speculative. Here we use super resolution microscopy to study the lateral organization as well as the functional relationship of plasma membrane-bound molecules involved in IAV infection. We find that molecules are organized in submicrometer nanodomains and, in combination with virus diffusion simulations, present a mechanistic view for how IAV first engages with AFs in the plasma membrane to then engage and trigger entry-associated membrane receptors.

## Introduction

Influenza A viruses (IAV) cause severe respiratory tract infections in humans often leading to seasonal local epidemics as well as periodic global pandemics [1]. During cell binding, IAV engages with low affinity attachment factors (AFs) as well as functional receptors to trigger cell entry by endocytosis. However, little is known about the lateral organization of both, AF and functional receptors, and how their organization translates into their functional role during virus infection.

The viral factor responsible for IAV-cell contact, the first step of infection, is the envelope protein hemagglutinin (HA), a trimeric glycoprotein that covers ∼ 90 % of the viral surface (Fig. 1A) [2]. The most common cellular AF for IAV is N-acetylneuraminic acid (also sialic acid), a highly abundant cell-surface glycan that within its glyosidic linkage can also encode IAV species specificity. Human-pathogenic IAV strains preferentially bind α-2.6-linked sialic acid, while avian-pathogenic viruses prefer to bind α-2.3-linked sialic acid. This specificity can be attributed to the topology of the glycan, which can more readily form contacts with receptor-binding domains of complementary HAs [3]. A common feature of glycan-protein interactions is their low affinity, which for HA-sialic acid lies in the millimolar range [4] and should make it challenging for the virus to form a stable interaction with cells. Although the glycan is highly abundant, which could lead to adhesion, single-virus tracking showed that the particles have some degree of freedom to explore the cell surface [5–7]. Indeed, it remains largely unclear how an initial low-affinity interaction, with particle mobility, can lead to a stable and specific virus-cell contact enabling a successful infection.

**Figure 1:**
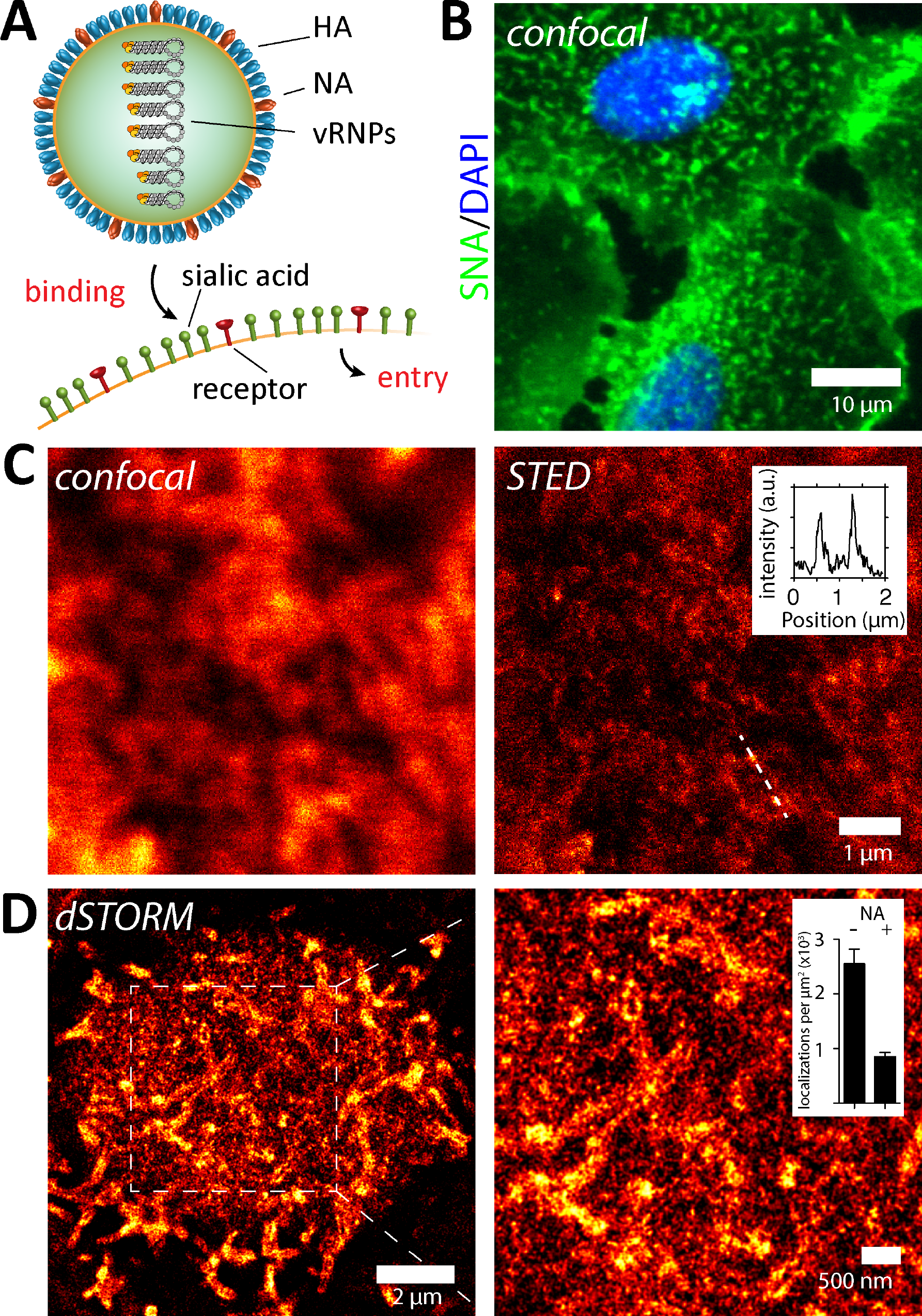
Sialic-acid containing IAV attachment factors are organized in nanodomains on A549 cells. (**A**) Influenza virus is an enveloped particle that encapsulates the segmented (-)vRNA genome built of 8 viral ribonucleoprotein particles (vRNPs). The viral membrane harbors the two glycoproteins hemagglutinin (HA) and neuraminidase (NA). HA is responsible for binding sialic acid (SA) containing attachment factors on the host cell plasma membrane. Upon cell-binding, the virus needs to activate functional receptors to trigger endocytosis. (**B**) Confocal imaging of live A549 cells labelled with SNA and Hoechst (DNA) feature a nonuniform SNA distribution across the plasma membrane. Large finger-like protrusions can be observed on the apical plasma membrane. (**C**) Confocal and STED live-cell imaging of A549 cells labelled with SNA confirms the existence of finger-like protrusions as well as a population of smaller nanodomains with diameter of ∼100 nm (**C**, right, inset). (**D**) Further, we utilized STORM imaging of A549 cells labelled with SNA. Reconstructed STORM images confirm two major structural features (1) finger-like protrusions as well as (2) small nanodomains. Cell treatment with neuraminidase (NA, 0.01 U/ml for 2h) led to a strong reduction of the localization density due to the cleavage and hence decrease local concentration of SA (**D**, right, inset).

After binding, IAV enters cells by receptor-mediated endocytosis, where clathrin-mediated endocytosis was shown to constitute the major [7], albeit not the only entry route [8]. Since the role of sialic acid (SA) as the primary AF is only passive and it cannot trigger endocytosis, an active signal-processing receptor must also engage to allow viral entry. Recently, receptor-tyrosine kinases were shown to be able to fulfill this function [9]. Specifically, it was shown that, among other receptor-tyrosine kinases, epidermal growth factor receptor (EGFR) was activated during and necessary for IAV cell entry. However, how IAV finds and activates EGFR has remained speculative. While molecular and structural information is available for both HA-SA [3] and EGFR-EGF interactions [10], much less in known about the spatial organization enabling EGFR activation during IAV cell infection.

Electron microscopy has provided a detailed picture of influenza viral particles [2] as well as its individual proteins [11]. However, imaging and quantitative analysis of cellular structures at the nanoscale remains challenging. Super-resolution microscopy represents an excellent tool to study the organization of cellular membranes at the scale of the viral particle (<100 nm) [12]. Here, we combined two complementary approaches, which together provide a versatile toolbox to study biological systems [12,13]. We used single molecule localization (SML) techniques known as stochastic optical reconstruction microscopy (STORM) and (fluorescence) photoactivated localization microscopy ((f)PALM) to image the organization of molecule in the cell membrane and track single molecules of EGFR [14–16]. We also applied stimulated emission depletion (STED) [17] to perform live-cell super-resolution microscopy.

We quantitatively analyzed the spatial organization of IAV AFs as well as EGFR on the surface of human alveolar epithelial cells. We found that AFs are organized in virus-sized clusters featuring a density gradient that decreases from the dense core to the periphery. Using these experimentally-determined characteristics, we investigated their role in virus-cell interactions and mobility with simulations. These simulations are in good agreement with virus tracking experiments, and together, they suggest that the spatial organization of AFs dominates virus-cell interaction during the early phase of virus infection. We further show that AF nanodomains overlap with EGFR clusters thereby enabling an AF-mediated EGFR activation. Interestingly, our results further suggest that pre-existing EGFR clusters are responsible for IAV-mediated receptor activation. We provide a novel view on the initial events of influenza virus infection and offer new insights into the functional role the spatial organization of cell surface AF and receptors.

## Results

### Sialic-acid containing IAV attachment factors are organized in nanodomains on the plasma membrane of A549 cells

To examine the spatial organization of IAV AFs within the plasma membrane of permissive epithelial cells, we labeled them with fluorescently-modified lectins. Specifically, we used the plant lectin *Sambuccus nigra* agglutinin (SNA), which selectively recognizes α-2,6-linked sialic acid moieties. As this specific sialic acid linkage is preferably recognized by human-pathogenic IAV, such as H3N2/X31 [4] used here, we used SNA as a primary IAV AF label (please see also supplementary note 1). Using confocal microscopy, we found that SNA strongly labelled the plasma membrane of live A549 cells (Fig. 1B), showing enrichment in finger-like protrusions that morphologically appeared to be microvilli. We then used STED microcopy to more carefully study the smoother regions of the plasma membrane between the microvilli. On live A549 cells, we detected a strong heterogeneity of SNA cell surface labelling including small clusters at the scale of 100 - 200 nm (Fig. 1C, right panel, inset).

Since small spherical H3N2/X31 virions have an average diameter of 120 nm [2], our next goal was to investigate the lateral organization of IAV AF at the scale of the virus-cell interface (radius < 60 nm). For this purpose, we imaged fixed A549 cells labelled with SNA conjugated to Alexa647 using STORM. STORM imaging confirmed our observations made with STED on live cells and revealed that SNA also labelled a variety of smaller structures that appeared on the flat parts of the plasma membrane (Fig. 1D). Such a heterogeneous plasma membrane carbohydrate distribution was also observed previously using Vero cells [18]. By labelling cells using antibodies against ezrin, an actin-binding protein that is highly enriched in microvilli [19], we confirmed that the larger structures are indeed microvilli (Fig. S2A). Microvilli, due to their narrow size, are not actively involved in endocytosis [20,21]; thus, we focused our quantitative analysis on the smaller AF cluster population.

### Quantitative analysis of SNA nanodomains

To describe the lateral arrangement of AF from our STORM data, we analyzed the distribution of localizations using an algorithm based on the detection of local density differences. To this end, we developed a cluster analysis routine (see *Methods*) that allowed us to extract geometrical properties of the clusters as well as an estimate of the number of AF molecules (Fig. 2). To identify and threshold the large microvilli cluster population, we first performed cluster identification on the ezrin localization maps (Fig. S2B). We found that the large ezrin clusters had dimensions of 10 - 50 nm across the short and 0.5 - 2 μm along the long axis (Fig. S2A). These parameters where then used to filter out the large cluster population corresponding to microvilli identified in AF localization maps. After filtering, we identified a heterogeneous population of small clusters with an average area of 0.016 μm^2^ (Fig. 2C). Since the cluster area was found to be at the same scale as the projected two-dimensional area of a spherical IAV (0.0079 μm^2^ for r = 50 nm), we took a closer look at the localization density within each individual cluster. For each localization, we identified the number of nearest neighbor localizations within a distance of three times the localization precision (3σ ∼ 30 nm) (Fig. S3). Interestingly, we found that AF clusters have an average 10-fold enrichment compared to the local background while some reach an even up to 20-fold increase in receptor density (Fig. 3). Using simulated AF domains, we observed that this local concentration effect can be partly mimicked by the localization precision (Fig. S4). However, in our experimental case this accounts only for an enrichment of < 8-fold (Fig. S4D).

**Figure 2:**
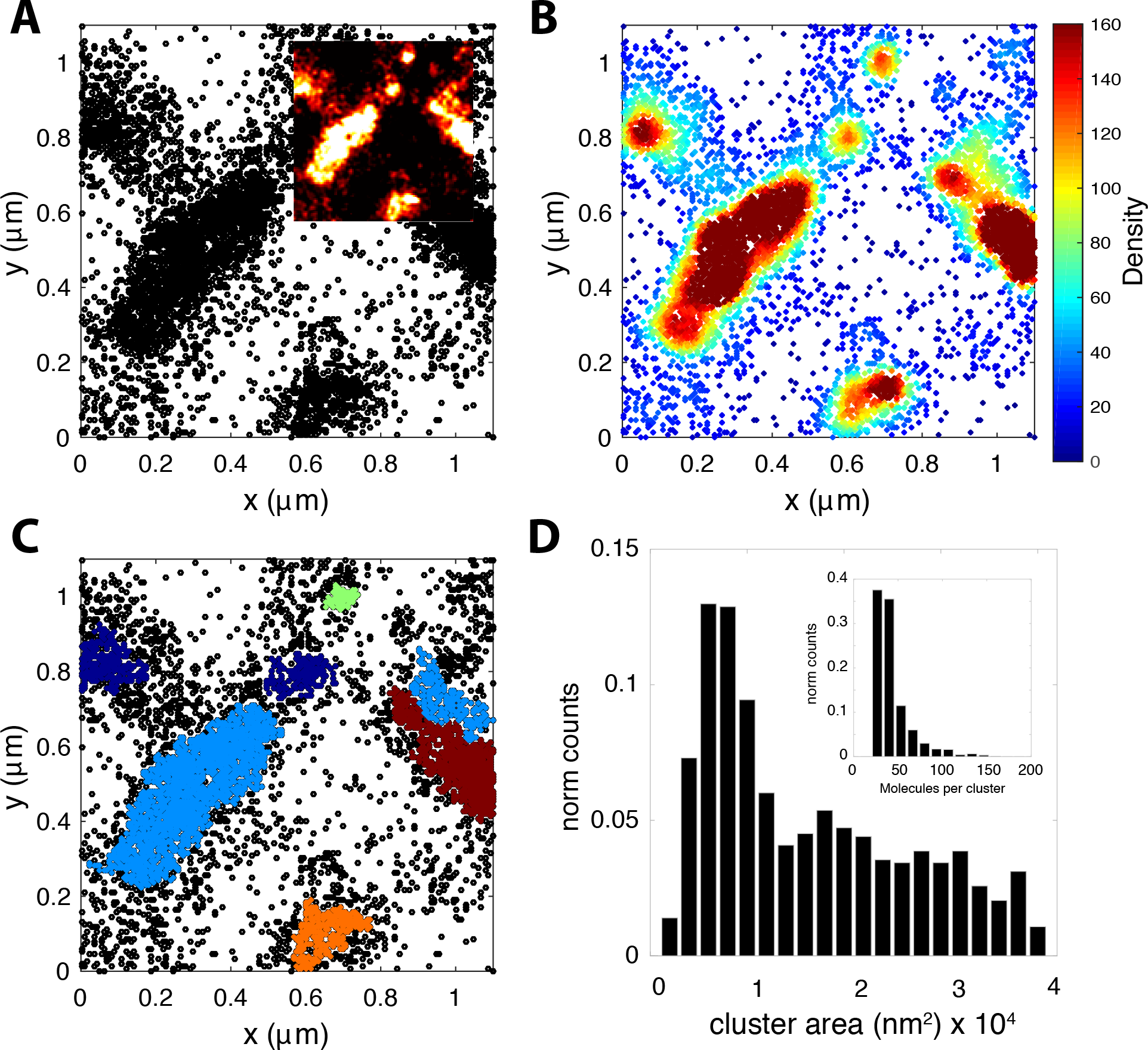
Density-based localization analysis reveals small SA clusters between microvilli. (**A**) Spatial distribution of STORM localizations from SNA-A647 on A549 cells showing the coexistence of two structural features, (1) large microvilli as well as (2) small nanodomains. The inset in **A** shows a rendered reconstruction (10 nm/pxl) of the localization map in **A**. (**B**) Density distribution of localizations shown in **A** within a search radius of 50 nm. Color scale according to number of neighbor localizations. (**C**) Final cluster identification with identified clusters in random color code. (**D**) Distribution of cluster area. The cluster identification allows quantification of the cluster area. After all identified clusters were filtered according to their size to selectively analyze non-microvilli structures, we found clusters with an area between 0.5 - 4 * 10^4^ nm^2^. Distribution of the number of molecules per cluster as estimated according to the number of localizations (**D**, inset).

**Figure 3:**
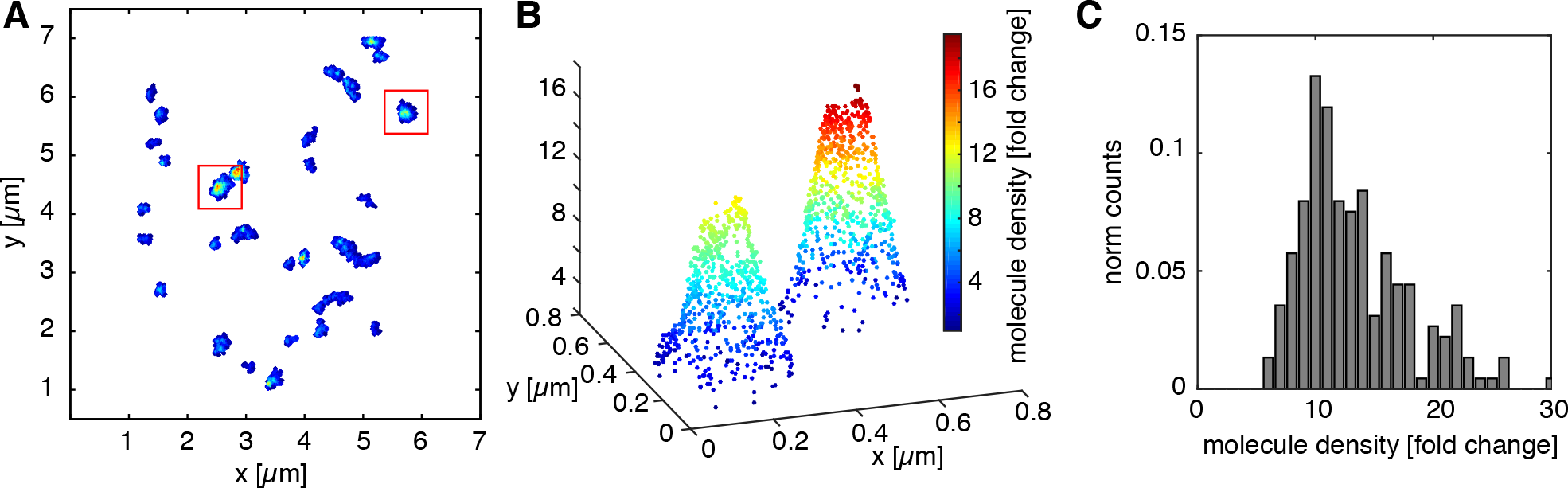
Small SNA clusters have an inner density gradient of localizations decreasing from the dense core to the cluster periphery. The inner structure of non-microvilli clusters was analyzed according to their local localization density. (**A**) Representative example of a membrane patch with identified clusters showing the inner density gradient. The color code represents the number of nearest neighbor localizations within a radius of 30 nm (i.e. the local localization density). (**B**) 3D plot of the two clusters boxed in **A**. The localization density is plotted on the vertical axis. (**C**) Distribution of the density difference between background and the cluster center over all identified clusters.

### IAV performs a receptor concentration-driven random walk on the plasma membrane

Our information on AF clustering led us to wonder how the overall IAV motion on the cell membrane would be affected by the heterogeneous local AF concentration. We established a simple diffusion model to simulate the behavior of individual viruses on the cell surface. The model assumes a two-dimensional random walk [22] where the virus undergoes periods of free diffusion (with diffusion coefficient *D_free_*) until it encounters a region of high AF concentration and becomes confined (with diffusion coefficient *D_conf_*). The time a simulated particle stays confined will depend on *D_conf_* as well as on the size of the confined region (i.e. the AF cluster size as measured using STORM). To identify and quantify confined regions, we establish a confinement index *I_conf_* denoting the probability for the particle to be confined at time *t* [23]. Even simulations of purely free diffusion will display periods of apparent confinement, due to the stochastic nature of thermal motion. We define a threshold level of confinement to exclude these random fluctuations, *I_thresh_* = 15, which allows identification of confined areas and comparison of the confinement dwell time (Fig. 4 A, D). Our simulations reveal a characteristic *I_conf_* signature of free (Fig. 4D) as well as confined diffusion (Fig. 4B and E and supplementary movie 1). In simulations of purely free diffusion, the confinement index does not rise above *I_thresh_*, which is in contrast to the case after including confinement zones (Fig. 4E).

**Figure 4:**
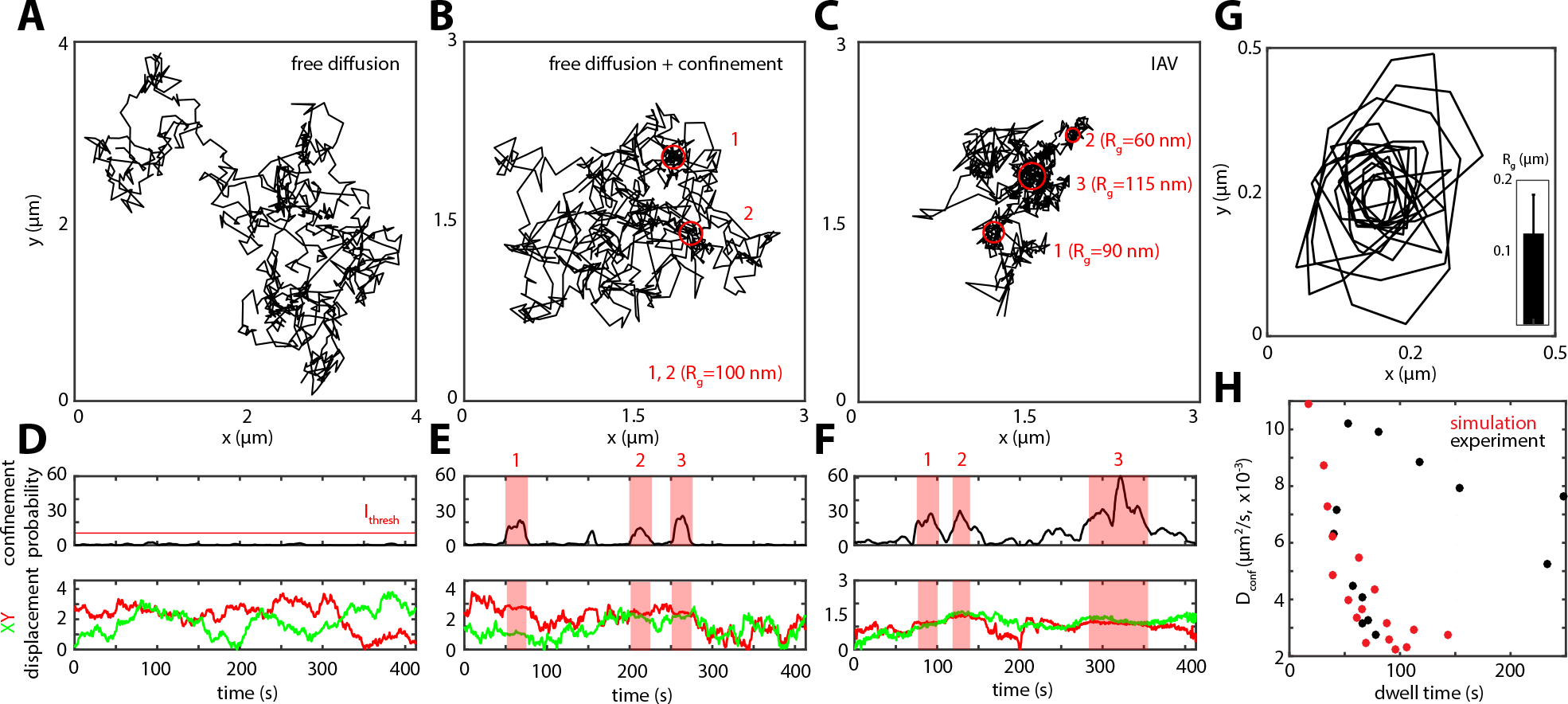
IAV performs a receptor concentration-driven random walk on the plasma membrane. Based on our quantitative analysis of the AF distribution on A549 cells, we hypothesize a motion behavior that is driven by the local AF concentration. We simulate this behavior initially as a 2D random walk with free diffusion coefficient D_free_ (**A**). (**B**) Next, we simulate AF clusters (red circles, r = 100 nm), which would due to the increased SA concentration lead to a temporal confinement (D_conf_ < D_free_). To identify confined regions within the simulated virus trajectories, we establish a confinement index I_conf_. Accordingly, a free diffusing particle shows only fluctuation in of I_conf_ (**D**), while the addition of temporal confinement leads to a clear increase that overlaps with stationary phases of the particle as visible in the XY displacement plot (**E**). We used the confinement probability to analyze experimental virus trajectories in particular the mixed type of trajectories (**C**) (see also Fig. S5) (**C**). I_conf_ shows a clear signature of temporal confinement (**F**) similar to the model prediction (**E**). As a further challenge for our model, we performed a subtrajectory analysis, thereby extracting the dwell time, Dconf as well as the area of the respective temporal confinement in our virus trajectories. (**G**) shows an overlay of the perimeters of the extracted confined regions as well as the average radius (R). From our simulated data, correlation of D_conf_ with the dwell time shows that a local increase in AF concentration (i.e. decreased diffusion) due to the encounter of an SA nanodomain leads to an increased local dwell time (**H**, red markers). We observed a similar behavior, when we tested the confinement dwell time in experimental virus trajectories (**H**, black markers).

To test whether our model is consistent with the behavior of IAV on the plasma membrane of living cells, we performed single-virus tracking on A549 cells. To this end, IAV was labeled with the lipid fluorophore DiD as described in *Methods*. Labelled viruses were diluted in infection medium (DMEM, 0.2% BSA) to a final concentration of 20 μg/ml (protein content) and viral aggregates were removed using a 0.2 μm sterile filter. We performed virus tracking at physiological conditions (37 °C) as well as conditions that suppress virus endocytosis (4 °C, dynasore treatment) to prolong the particles’ residence time on the cell surface. During trajectory analysis, we observed different modes of movement, which we classified into four types: (1) confined, (2) ballistic, (3) drifting and (4) mixed (Fig. S5). The ballistic movement is directional, and goes up to speeds of 1-2 μm/sec; thus, we assigned it to microtubule-associated transport as previously described [6,7]. Since this type of movement follows a successful virus internalization, we expected to see it decrease in frequency after blocking endocytosis. Indeed, the fraction of ballistic trajectories dropped from 30% to below 5% when we imaged at low temperature or in the presence of 40 μM dynasore, a dynamin and thus clathrin-mediated endocytosis inhibitor [24]. Interestingly, we observed a marked increase of the other three motion classes, supporting the idea that they take place at the plasma membrane (Fig. S5). When we took a closer look at the mixed class of trajectories, we found regions of extended IAV residence time indicating spatial confinement alternating with free diffusion (Fig. 4C). Consequently, we applied our confinement analysis to the mixed IAV trajectory class. Interestingly, our trajectory analysis could detect pronounced areas of confinement that indeed alternate with regions of free diffusion (Fig. 4F). Hence, our data is consistent with our model of an AF concentration-driven cell surface motion.

If AF islands of different lateral concentrations coexist in the plasma membrane, as observed using STORM, these regions could serve as binding platforms for diffusing viruses. According to our simple diffusion model, we assume that the local AF concentration dominates *D_conf_* (see *Methods*), which in turn determines the particles dwell time inside the confined regions. Hence, to test if *D_conf_* correlates with the dwell time, we performed a sub-trajectory analysis on experimental trajectories, where each trajectory was screened for temporal confinement according to *I_conf_*. For each confined region, we then identify the dwell time as well as *D_conf_* (see *Methods*). We observed indeed that the dwell time is negatively proportional to *D_conf_* as predicted by our diffusion model (Fig. 4H). Subtrajectory analysis further allowed us to estimate the spatial dimensions of the confined regions (Fig. 4G). We found an average radius of 104 nm corresponding to a median area of 5.7*10^4^ nm^2^, which is only slightly larger than the cluster size found using STORM (Fig. 2D). Together, we link the lateral organization of AFs, characterized using STORM, with live-cell IAV tracking data. The structural information served as an input for a diffusion model, whose predictions are consistent with dynamic single IAV plasma membrane motion. Our results suggest that IAV-cell binding and its dynamic surface movement are dominated by the local AF concentration.

### EGFR is organized in nanoclusters that overlap with SNA domains

Since sialic acid cannot transmit a signal into the cell and thus only serves as an AF for the virus, we wondered about its organization relative to that of the functional receptor EGFR. We investigated the lateral organization of EGFR in A549 cells using fluorescently labelled anti-EGFR antibodies. In our infection experiments, the cells were pre-incubated in serum-free infection medium (30 min) before viruses where added, a common procedure for influenza virus infection. To reproduce these conditions, we also performed a serum-free pre-incubation before the cells were fixed and immunolabelled for EGFR.

Using STORM imaging, we found that EGFR is present at much lower concentration on the cell surface compared to SA-conjugated AF, but, interestingly, also localized in nanodomains (Fig. 5A). Due to the sparsity of EGFR labeling, which might lead to false detection of protein clusters due to fluorophore blinking [25], we established an adapted analysis scheme based on the photophysical characterization of the fluorescent probe used in our experiments. As shown in Fig. S6, by imaging sparsely spread isolated molecules, we first measure the dark time *t_D_* as well as the spread of localizations originating from a single molecule ***Δ**(x,y)* (i.e the localization precision σ). Both parameters can then be used to correct the localization maps for blinking and allow a more accurate cluster identification as well as estimation of the number of molecules (Fig. S6).

**Figure 5:**
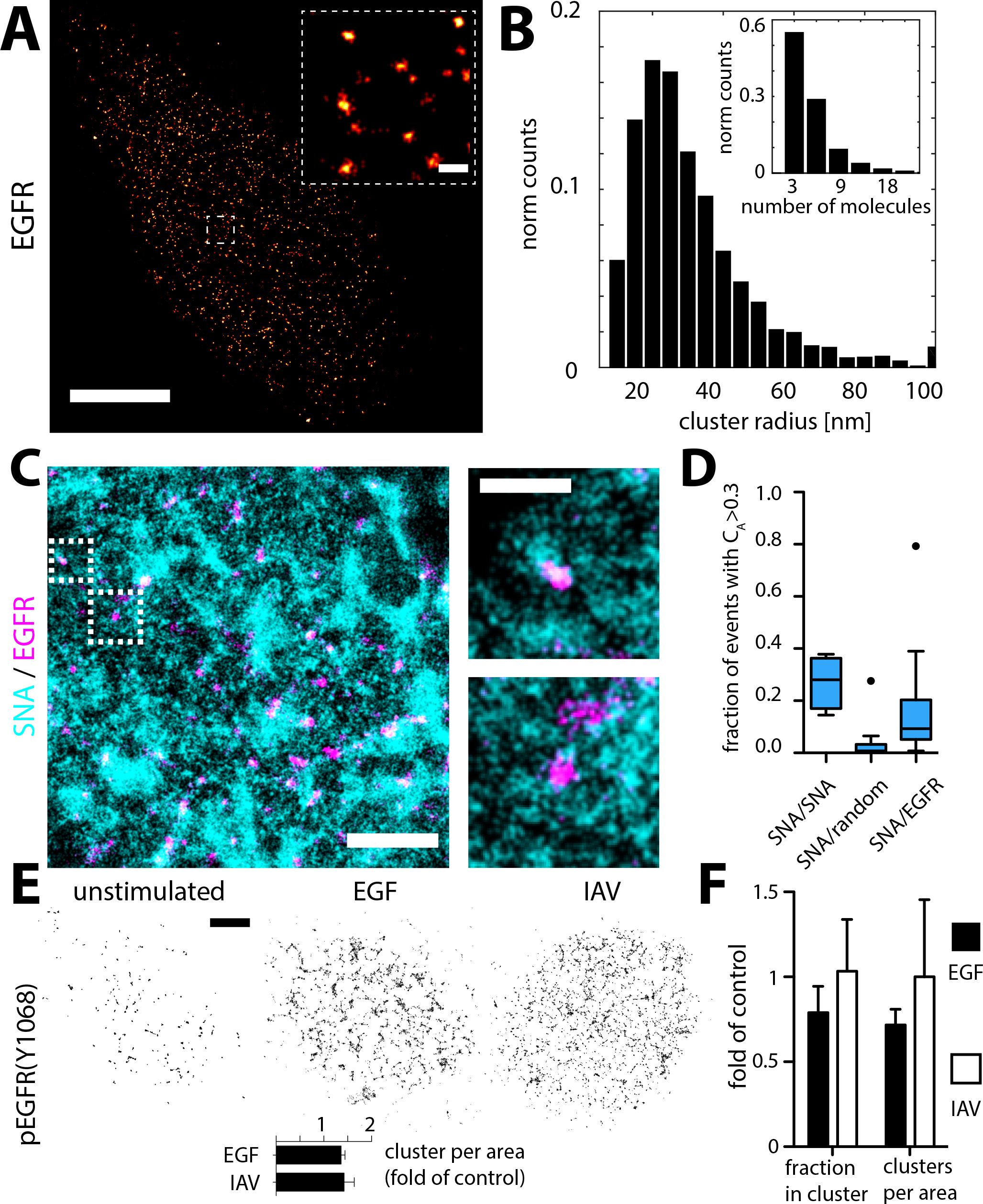
EGFR is organized in nanodomains that overlap with SNA domains. (**A**) A549 cells were labelled with antibodies against EGFR. STORM imaging revealed a clustered organization of EGFR on the apical plasma membrane. The clusters have an average diameter of 60 nm and contain about 5 - 10 molecules (**B**). Scale bar 1 μm. Inset: 100 nm. (**C**) Two-color STORM imaging of A549 cells labelled with SNA and anti-EGFR antibodies. The two panels on the right show larger magnification of the boxed areas in the left panel. Scale bars: 500 nm (left panel), 200 nm (right panel). The degree of colocalization was quantified using coordinate-based colocalization, where each localization is associated with a colocalization value C_A_. (**D**) Box plots of C_A_ distribution of SNA localizations when colocalized with (1) SNA, (2) a random distribution of localizations at equal density as EGFR and (3) EGFR. After stimulating the cells, we found that phosphorylated EGFR (Y1068) is also localized in nanodomains, suggesting activation of pre-formed cluster. Although a small population of clusters seems to be phosphorylated without stimulus, we observed an increase in the activated cluster population after stimulation with IAV or EGF (**E**, lower panel). To test for a potential redistribution of EGFR, we looked at the entire population after stimulation. While after EGF stimulation, we could observe a reduction of the clustered protein fraction as well as the cluster density per area, we could not detect such a protein redistribution after IAV stimulation (**F**).

After blink correction and cluster identification, we found that in the absence of EGF stimulation between 30 - 60 % of the EGFR molecules reside in clusters. The clusters have an average diameter of 29 nm and are composed of on average 6 molecules (Fig. 5B), which is in agreement with previous studies using electron microcopy [26] and FRET [27]. Since IAV directly binds to sialic acid on the cell surface, one way to facilitate IAV-EGFR interactions would be for EGFR and AF to occupy the same regions on the plasma membrane. To test this hypothesis, we performed two-color STORM imaging (Fig. 5C) using A549 cells co-labeled for AF (with SNA) and EGFR. Indeed, we found that EGFR clusters overlap with SNA-labeled membrane domains. However, since sialic acid AFs are much more abundant than EGFR, their colocalization could occur simply by chance. To examine this possibility, we performed a quantification based on coordinate-based colocalization (CBC) [28]. CBC analysis provides an estimate for the spatial correlation of two localization datasets, reflected in the colocalization parameter *C_A_*. To get a better indication about the extent of colocalization in our SNA/EGFR dataset, we added two controls to our analysis. We performed an experimental positive control by using the lectin SNA conjugated with two different fluorophores (denoted SNA/SNA). As a negative control and to take the difference of localization density into account, for each two-color field of view in SNA/EGFR, we simulated a random dataset at the same density as the EGFR dataset (denoted SNA/random). Finally, we counted localizations with C_A>0.3_ as colocalized (Fig. S7). As shown in Fig. 5D, the negative control SNA/random reaches with C_A>0.3_= 0.035 the lowest level of colocalization only accounting for random colocalization, while the experimental positive control SNA/SNA reaches the highest score of C_A>0.3_ = 0.27. CBC analysis of SNA/EGFR colocalization reached C_A>0.3_ = 0.17, suggesting that EGFR and SNA do not randomly colocalize, but indeed share the same membrane compartments.

If an infecting IAV encounters an EGFR cluster following attachment to sialic acid AFs, we would expect EGFR activation upon IAV adsorption. Next, we wanted to test whether the cell’s EGFR pool responds to stimulation using an antibody that recognizes a phosphorylated tyrosine (Y1068, pEGFR) previously shown to be involved in IAV-induced EGFR activation [9]. Interestingly, we found a fraction of pEGFR nanodomains even under unstimulated conditions. However, following stimulation with both, EGF or IAV, we observed an increased number of pEGFR clusters per area on the plasma membrane (Fig. 5E). Notably, in order to keep the signal at the plasma membrane, endocytosis was slowed down by stimulating the cells on ice.

In order to better understand how IAV binding leads to EGFR activation, we took a closer look at the properties of individual EGFR clusters. It was previously hypothesized that IAV binding leads to a local concentration of EGFR proteins in plasma membrane clusters eventually leading to signal activation [9], an effect that was also observed before in BHK cells upon EGF stimulation [29]. To test this hypothesis, we labeled unstimulated as well as IAV-adsorbed cells using anti-EGFR antibodies. After EGF stimulation, we observed a decrease in the clustered molecule population as well as the number of clusters per area (both by on average 20%) (Fig. 5F). Surprisingly, we did not find evidence for a significant redistribution of EGFR after IAV-cell binding (Fig. 5F). In addition, we could not detect an effect on the cluster size or the number of molecules in a cluster (Fig. S8).

### EGFR forms long-lived nanodomains in living cells

Our results indicate that pre-formed receptor clusters might be involved in IAV-induced EGFR activation. While such a mechanism was previously hypothesized [30], it was never shown that receptor clusters can reach lifetimes in the plasma membrane that would allow this type of activation. To estimate the lifetime of EGFR clusters, we turned to live-cell microscopy experiments using A549 cells expressing EGFR tagged with the photo-convertible protein mEos3 [31]. We used an EGFR variant that was previously shown to be fully functional in a mammalian cell expression system [32]. Photoactivation of only a small subset of EGFR-mEos3.2 molecules allowed us to localize individual molecules which can then be tracked over consecutive frames, and renewed by further photoactivation (single-particle tracking, sptPALM) [33]. We performed sptPALM on the apical as well as the basolateral plasma membrane of live A549 cells in the absence of EGF, resulting in high-density protein diffusion mapping (Fig. 6). Calculation of the mean-squared displacement (MSD) as a function of the lag time ***Δ**t* allowed us to determine the instantaneous diffusion coefficient *D* along the trajectory. We found a broad range of diffusion coefficients ranging between 1 and 10^−3^ μm^2^/sec (Fig. S9), indicating that mobile and immobile protein fractions coexist in the plasma membrane. Indeed, after classifying trajectories based on their diffusion coefficient *D*, MSD vs. **Δ***t* plots revealed a rather linear relationship for trajectories with *D* > 0.05 μm^2^/sec, while for trajectories with *D* < 0.05 μm^2^/sec the curve approached a maximum at large **Δ**t values (Fig. S9). This characteristic time dependence of the MSD vs. **Δ***t* curve also indicates mobile free diffusing proteins co-existing with immobile or confined-diffusing EGFR proteins in the plasma membrane.

**Figure 6:**
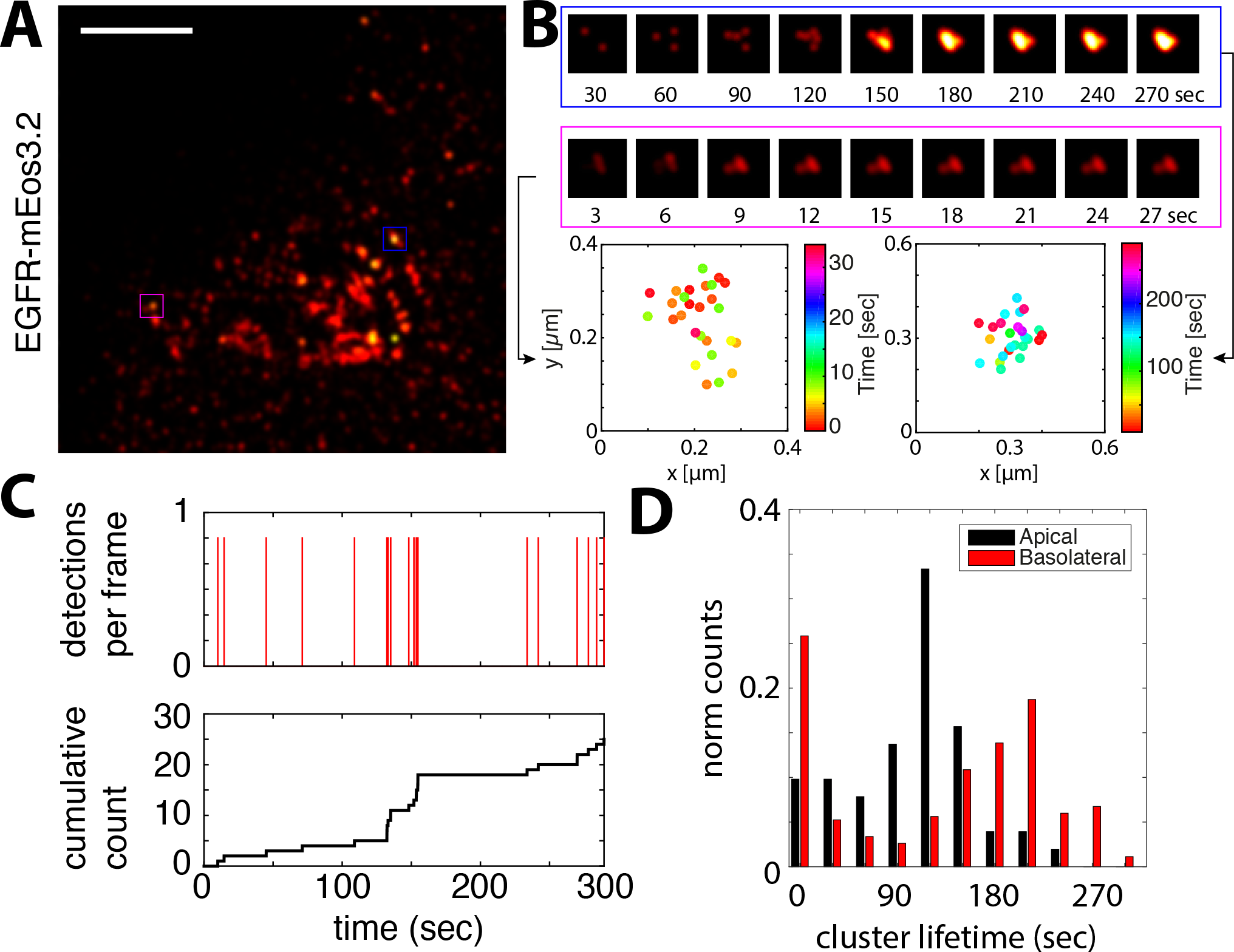
Live-cell super-resolution imaging reveals long-lived EGFR clusters in living cells. EGFR coupled to the photo-convertible protein mEos3 was expressed in A549 cells. Subsequent PALM imaging allows to study EGFR distribution in live cells at the single protein level. In the absence of any stimulus, we could detect nanodomains of EGFR within the apical and also the basolateral plasma membrane (**A**). Scale bar: left panel, 1 μm. The image in **A** shows a maximum projected map of single molecule localizations recorded over a period of 10 min. **B** shows two cluster examples as a cumulative density distribution (upper panel) as well as XY scatter with the colorscale according to time at which the localization was detected (lower panel). While the projection of all localization allows to identify protein clusters, we can use the time information to further estimate the cluster lifetime. As shown in **C**, cumulative counting of individual localizations within a clustered region gives direct information of the minimum cluster lifetime. **D** shows the corresponding lifetime distribution of EGFR clusters recorded at the apical as well as the basolateral membrane in the absence of any stimulus.

The same data was then used to construct a map of all detected localizations, which revealed a non-homogeneous distribution, with proteins appearing clustered in nanodomains (Fig.6A). Using our custom cluster identification (as in Fig. 2), we found EGFR clusters with a diameter ranging between 30 and 300 nm, thereby confirming the clustered organization of EGFR in living cells as seen after cell fixation using STORM. In addition, because the cells were alive, we could use the time-resolved single molecule detection to quantify the temporal stability of EGFR clusters (Fig. 6B). Such an approach was used previously to quantify polymerase 2 clustering in live cells and is referred to as time-correlated PALM (tcPALM) [34]. We selected only regions identified using density-based clustering to perform cumulative localization counting (Fig. 6B, C). We obtained a distribution of EGFR cluster lifetimes with an average of 140 seconds (Fig. 6D, apical). Notably, this lifetime only provides a lower estimate since the cluster might have existed before starting the acquisition and might also be present after the last cluster molecule is bleached.

## Discussion

Understanding the initial phase of virus infection is crucial for the development of effective countermeasures such as adhesion inhibitors that catch viral particles before they can engage in the first virus-cell contact [35,36]. After successful binding of SA-containing AF on the cell surface, the virus has to find its functional receptor to enter and infect its target cell. While these two steps are crucial for IAV infection, it remained largely speculative how the virus finds a way to efficiently bind a cell and engage with a functional receptor. Our results provide a quantitative structural view of the lateral organization of virus AF and receptors while suggesting a functional link between cell binding and receptor activation. Several studies have tracked fluorescently-labeled IAV on and inside respective target cells [5–7]. It was found that IAV particles after forming the initial cell contact move in an actin-dependent way for on average 6 min before endocytosis [6]. In a later study, using multicolor imaging of virus and clathrin, this period was assigned to be important for the induction of clathrin-mediated endocytosis [7]. However, how these processes are linked and what are the structural determinants remained unclear.

We used super-resolution microscopy to investigate the structural organization of AF as well as one functional receptor - EGFR - at the scale of the infecting virus. We show that SA-conjugated AFs are organized in a heterogeneous population of differently sized nanodomains. We find that one labeled structure can be assigned to microvilli, a dominant topological feature of the apical cell surface in epithelial cells and further focused our attention on the non-microvilli cluster population found on the flat and endocytosis-active regions of the plasma membrane. Here we found smaller nanodomains ranging in diameter from about 50 - 300 nm. Interestingly, when we looked at the local localization density per cluster, we found that the clusters differ in their molecular density where some show an up to 20-fold increase in molecule concentration towards the cluster center. To test if the grouping of AF in dense clusters is advantageous for virus binding, we constructed a simple binding simulation (Fig. S10). In one part of the simulation, we gradually shift AF molecules from a random position into a clustered organization, while the total number of molecules stays constant. In a second approach, we simulate a stable population of clusters within a background of free individual molecules and vary the number of molecules per cluster. In both cases, we simulate a spherical virus particle and project its size as a landing spot on the simulated molecule surface. Successful binding is counted if the virus can at least bind 10 molecules. As shown in Fig. S10, we find a strong positive correlation between clustering and receptor binding clearly indicating that nanoclustering enhances the probability of efficient virus binding. In addition, as it was also suggested before for DC-SIGN [37], this heterogeneous cluster organization might also broaden the binding capability of the cell surface and effectively provide the virus particle with a range of binding times to explore the cell surface. We hypothesize this behavior and simulate viral movement based on the availability of AF resulting in a predicted random walk motion that is intercepted by temporal confinement due to local AF enrichment. We went on to test if this behavior can be observed experimentally. Our single-virus tracking experiments showed four major types of movement. While fast directed transport was shown to be microtubule-associated inside the cells [6], slow drifts, confinement and diffusive motions are characteristic for plasma membrane movement [38]. Indeed, when we used conditions of hindered endocytosis (i.e low temperature or dynamin inhibition), we detected an increase in the slow and almost complete disappearance of the fast trajectory types. Interestingly, we also found that the dwell time in confined areas during diffusive motions correlates with the confinement diffusion coefficient *D_conf_*. This follows our simulation confirming our basic hypothesis that virus plasma membrane motion is dominated by the surface concentration of available AFs. We provide a functional link between the clustered organization of virus AF and their role for virus-cell binding.

Having formed a stable interaction with its host cell, IAV enters the cell by endocytosis. EGFR and other receptor tyrosine kinases were shown to be activated during and promote IAV-cell entry [9]. It was suggested that IAV binding leads to EGFR clustering and the formation of an active signaling platform [9]. We found that also under unstimulated conditions, EGFR is mainly (up to 60 %) localized in small nanodomains with a mean diameter of 29 nm containing on average 6 molecules. In the canonical activation model, EGFR binds its substrate EGF, undergoes dimerization and subsequent autophosphorylation, thereby inducing a variety of signaling cascades [39]. However, an additional level of higher oligomeric EGFR clusters has been shown across different cell types. Their reported diameter ranges from 50 nm [29] over 100 - 300 nm [26,30,40] up to near micrometer sizes [41], with molecule numbers between <10 [42] up to thousands [41]. Clustering and cluster activation of EGFR was suggested to facilitate receptor activation which might play a role in tumor development [30]. Further, the EGFR cluster size was shown to respond to activation [29] suggesting a lateral molecule redistribution. Upon binding of EGF or IAV, we probed the cells with antibodies specifically detecting the autophosphorylation site Y1068, shown to be involved in IAV-mediated EGFR activation [9]. We found an increased signal at the plasma membrane which, when imaged in STORM, was found concentrated in nanodomains. Interestingly, the tetrameric SNA could not activate EGFR suggesting that IAVs higher multivalency is needed for efficient receptor activation (Fig. S11). At this point, we hypothesized two scenarios in which, during activation, EGFR either (1) assembles into activated clusters or (2) pre-existing clusters become activated. Hence, we tested the organization of EGFR under stimulated conditions. While we observed a reduction of the clustered molecule fraction as well as the number of clusters per area upon EGF stimulation, an effect that was observed previously for Erb2 [43], we could not detect a major redistribution in response to IAV attachment. Also, using STORM, we could not detect a change in cluster size and molecule composition following either stimulation (Fig. S8). We conclude that intercluster spatial rearrangements below our resolution eventually lead to cluster activation. Methods that are more sensitive to protein-protein distances below 20 nm such as FRET could be used to test this hypothesis. Then to test if EGFR clusters exist long enough to allow their activation, we conducted live-cell sptPALM. Our PALM imaging could confirm the existence of nanodomains within the plasma membrane of unstimulated cells. Consequently, we performed spatial clustering to find zones of EGFR enrichment and measure their lifetime. We found that the lifetime of EGFR nanodomains in both, the apical as well as the basolateral membrane went up to 2 to 4 minutes. While such a long cluster lifetime allows activation of pre-existing clusters, this observation raises the question for the stabilization of EGFR clusters. Hence, we tested the stability of EGFR clusters upon treatment with classical membrane domain-destabilizing conditions such as actin depolymerization (latrunculin A) and cholesterol extraction (methyl-β-cyclodextrin). As also observed before [29,30], our results suggest cluster destabilization following both perturbations (Fig. S11B) indicated by an increased fraction of unclustered EGFR molecules. Interestingly, we found a stronger effect after cholesterol depletion, a condition that was previously shown to attenuate IAV replication [9]. Very long receptor cluster lifetimes were observed previously for class I major histocompatibility complex (MHC) molecules, that form actin-stabilized domains [44,45]. However, one can speculate about the function of membrane receptor clusters [46]. It can be excitatory as shown for T-cell receptor [47] and linker for activation of T-cell (Lat) [48,49] as well as LFA-1 [50] or CD36 [51]. These nanodomains render the cell highly sensitive to small amounts of signaling molecules and due to the high local receptor concentration allow a fast signaling response [39]. Such a function seems likely for EGFR clusters involved in IAV cell entry observed in our study. But their function can also be inhibitory as shown for the B-cell receptor [52] or its negative co-receptor CD22 [53]. Receptor nanodomains could even engage in modulating the signaling output. As EGFR sits at the top of a broad array of signaling cascades, an asymmetric distribution of receptors could enable cells to rapidly respond and process different stimuli [39].

In summary (see Fig. 7), our results show the compartmentalization of the cellular plasma membrane in A549 cells. We found that both of the primary IAV-binding molecules, the AF sialic acid as well as the functional receptor EGFR, are organized in nanodomains. We further build a functional link between the lateral membrane organization and its impact on virus infection. While AFs forms dense clusters, it provides a multivalent binding platform allowing stable virus attachment. The diversity of those clusters (i.e. their size and molecule concentration) results in a spectrum of binding times. In our case, we observed dwell times between 5 - 20 sec for mixed diffusive motion or much longer (several minutes) if we include confined trajectories. Since EGFR clusters overlap with SA-enriched areas, IAV can indeed reach a functional receptor, while diffusing between AF islands (Fig. 7). Finally, our results suggest that pre-existing EGFR clusters become activated during virus binding. Quantitative super-resolution microscopy has provided a versatile toolbox to study the lateral organization of the plasma membrane to understand its structure-function relationship. Our study provides a first example for how membrane compartmentalization can engage in and modulate IAV cell binding and receptor activation.

**Figure 7:**
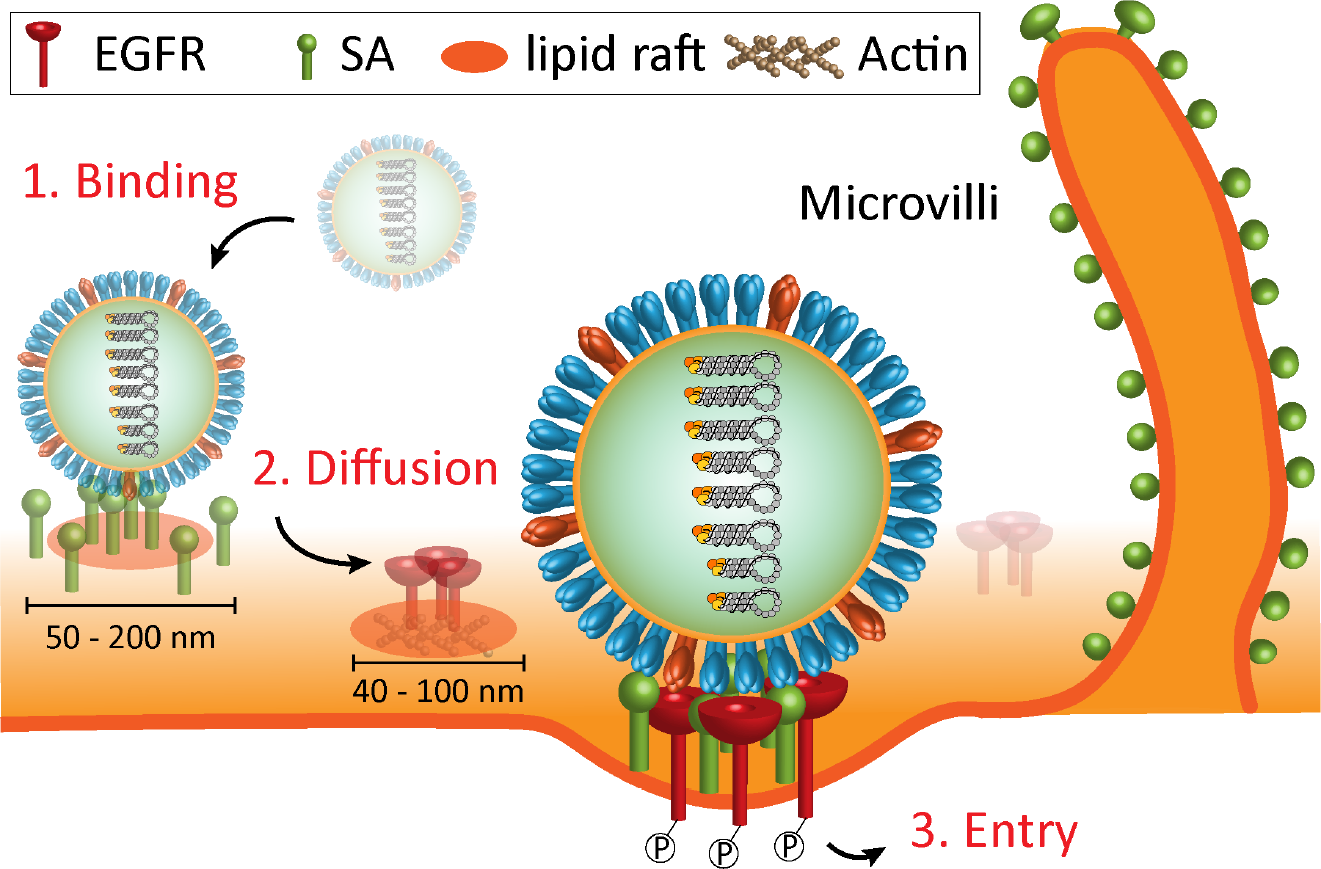
Model for IAV-mediated cell binding, receptor search and activation. Using quantitative STORM imaging, we could show that SA-conjugated IAV AF as well as one functional receptor, EGFR, form nanodomains in the plasma membrane of A549 cells. While dense AF nanodomains constitute an attractive multivalent binding platform, their diversity in local AF concentration suggests a variety of different residence times for which IAV would stay bound within these domains. Using single-virus tracking, we observed a mixed diffusive - confined motion, that could be simulated using our quantitative SA cluster information. These data suggest a receptor concentration-driven lateral search mechanism between SA enriched nanodomains. Eventually, since AF domains partly overlap with EGFR, IAV encounters a functional receptor that can be activated to signal cell entry. Our data further suggest that a stable pre-formed EGFR cluster population is activated during IAV stimulation, thereby possibly facilitating efficient signal transduction. EGFR clusters are stabilized by lipid rafts as well as cortical actin.

## Materials and Methods

### Ethics statement

Work with embryonated chicken eggs was conducted in the lab of Prof. Andreas Herrmann (Institute for Biology, Humbold-Universität zu Berlin, Germany) in accordance with European regulations and approved by the Berlin state authority, Landesamt für Gesundheit und Soziales. Influenza A (H3N2) X-31 was propagated in the allantoic cavities of 10-day old embryonated chicken eggs (Lohmann Tierzucht GmbH, Germany) as described previously [54].

### Cells and Viruses

A549 cells (ATCC CCL-185) were kindly provided by Dr. Thorsten Wolff (Robert-Koch Institute Berlin, Germany). A549 cells were cultured in Dulbeccos Modified Eagles Medium (DMEM), supplemented with 10 % fetal calf serum (FCS). The cells were passaged every 3-4 days. One day prior to the experiment, the cells were detached from the cell culture flask using 0.5 % Trypsin/EDTA for about 10 min. The cells were diluted in fresh DMEM and 3*10^5^ cells were seeded on fibronectin-coated 25 mm round glass slides (Menzel, # 1.5). Influenza A (H3N2) X-31 was propagated in the allantoic cavities of 10-day old chicken eggs (Lohmann Tierzucht GmbH, Germany) as described previously [54]. Purified viruses were stored at −80 °C. Virus aliquots were thawed on the day of the experiment and kept on ice until further use. All chemicals if not otherwise stated where purchased from Sigma-Aldrich. Cell culture media were purchased from Life Technologies.

### A549 cell infection

One day prior to the experiment, 3*10^5^ cells were seeded on fibronectin-coated 25 mm round glass slides. For infection experiments (Fig. S1), the cells where either incubated in serum-free medium for 30 min (control) or in DMEM supplemented with 100 ng/ml human EGF (R&D Systems) for 90 min to remove EGFR from the cell surface. Cells were infected with IAV X-31 (MOI ∼1) in infection medium (DMEM, 0.2 % bovine serum albumine (BSA)) for 30 min before the medium was changed and the cells were further incubated for 5 h in infection medium. The cells were washed in pre-warmed PBS and fixed in freshly prepared 4 % PFA (Alpha Aesar) for 10 min at room temperature. After a 25 min fixation/blocking step in PBS supplemented with 0.2 % Triton X-100 and 0.2 % BSA, the cells were incubated with the primary antibody (anti influenza nucleoprotein (NP), Millipore) for 1 h. The cells were washed three times 10 min in PBS before further incubated with secondary antibodies (goat anti-mouse, Alexa 555 conjugate, Life Technologies) for 1 h. Finally, the cells were washed in PBS, stained with DAPI (0,2 μg/ml in PBS for 10 min) and mounted on standard microscope slides with Mowiol (Carl Roth). The slides where imaged using a Zeiss Axioplan epifluorescence microscope. Ten overview images (20 x magnification) were acquired for each condition and nuclear NP signal was quantified using Cellprofiler[55].

### Single-virus tracking

IAV H3N2/X-31 were incubated with 50 μM of the lipid dye DiD (Life Technologies) for 2 h at RT. To remove the free dye, viruses were either pelleted (50.000 g for 5 min) or purified using a NAP5 size exclusion column (GE Healthcare). Immediately before the experiment, virus aggregates were removed using a 0.2 μm pore size filter. Labeled viruses were added to A549 cells grown in 35 mm poly-L-lysine coated glass bottom petri dishes (MatTek Corp.) and allowed to attach on ice for 10 min. The cells were washed with PBS and overlaid with 2 ml pre-warmed, serum and phenol red-free DMEM supplemented with 100 mM Hepes. Unless otherwise stated, the cells were kept either at 4 or 37 deg throughout the experiment. For the perturbation experiments, the cells were pre-incubated in DMEM supplemented with 50 μM nocodazole (Sigma) or 40 μM dynasore (Sigma) for 30 min. The drugs were kept present throughout the experiment. Low temperature incubation was achieved using a custom build microscope temperature chamber. DiD was excited with 633 nm laser light, which was reflected on the sample by a 488/633 nm dichroic mirror. Emission light was collected using a 60x PlanApo VC oil-immersion objective (Nikon) and imaged onto an EMCCD camera (Andor iXon, Andor Technology). Images were recorded at 2 frames per second for 10 min. Image stacks were processed and the trajectories were build using ParticleTracker for ImageJ [56]. The trajectories were further analyzed using custom MatLab (Mathworks) scripts. To identify and characterize temporal particle confinement, we used the method developed by Simson *et al*. [23] implemented into our custom analysis pipeline. Briefly, the algorithm takes a segment of the trajectory and determines if the particle moved according to a given free diffusion coefficient (D_free_) within the segment, i.e. if the particles stays in a predicted region. This is translated into a confinement probability/index I_conf_. Since the identification depends on the length of the segment *S* [23], we optimized S using simulated trajectories resulting in S = 5 s for our analysis.

### Trajectory analysis and single particle tracking simulations

Random brownian particle trajectories were generated using the script package *msdanalzer* [57] incorporated into a custom MatLab routine. Single virus trajectories were analyzed as described above. Trajectory and sub-trajectory analysis was performed using *msdanalzer* to retrieve diffusion coefficients from mean square displacement (MSD, <r^2^>) vs. lag-time (Δt) plots. MSD vs. lag-time plots were fitted according to the type of motional behavior, which was either free diffusion (<r^2^>= 4D_free_Δt) or, for sub-trajectory analysis, confined (sub)diffusion (<r^2^> = <r^2^> (1-A_1exp_ (−4A_2_D_conf_Δt/<r2>)). We found for IAV a mean free diffusion coefficient D_free_ = 0.041 μm^2^/s.

Using D_free_ as well as a time step (Δt = 0.5 s), the displacement of a freely diffusing particle follows a Gaussian distribution with standard deviation given by σ = sqrt(4D_free_Δt). Temporal confinement was introduced by generating a sub-trajectory using D_conf_. For the dwell time simulation, we generated random trajectories that run into a confinement region characterized by D_conf_ with a diameter of 50-300 nm according to the size of SNA clusters from STORM measurements (see also supplementary video 1). Confined regions were identified using the confinement index I_conf_ and the time the particle spends confined with I_conf_ > threshold was taken as the dwell time. D_conf_ was varied as shown in Fig. 2.

### Preparation of labelled lectins and antibodies

Unconjugated *Sambuccus nigra* agglutinin (SNA, VectorLabs) or anti-EGFR antibodies (Sigma) were diluted to 0.6 mg/ml in 100 µl PBS (supplemented with 50 mM NaHCO_3_). AlexaFluor 647 NHS ester (Life Technologies) or Star Red NHS ester (Abberior) was added at a final concentration of (150 μM) and the solution was incubated for 30 min at room temperature. 100 μl PBS were added and the solution was applied to a NAP5 size exclusion column (GE Healthcare) pre-equilibrated with PBS. 300 μl fractions were collected in a 96-well plate and analyzed by ultraviolet - visible spectroscopy (Nanodrop2000, ThermoFisher). Peak protein fractions were collected and the degree of labelling calculated. The labelled lectin and antibody fractions were stored at 4 °C until further use.

### SMLM sample preparation

One day prior to the experiment, 3*10^5^ A549 cells were seeded on fibronectin-coated 25 mm round glass slides. For SNA imaging, the cells were washed in pre-warmed PBS and fixed for 10 min in freshly prepared 4 % paraformaldehyde (Alpha Aesar). The cells were blocked in blocking solution (5 % BSA in PBS) and incubated in 50 μg/ml SNA diluted in blocking buffer for 30 min. The cells were washed 3 times in PBS and post-fixed in freshly prepared 4 % paraformaldehyde for 10 min at RT. For EGFR labelling, the cells were washes, fixed and blocked as described above then incubated with anti-EGFR primary antibodies conjugated to Alexa 647. For two-color imaging, the cells were incubated with unconjugated primary anti-EGFR antibodies for 1h at RT. The cells were washed three times in PBS and further incubated with a solution of 5 μg/ml Alexa 750-conjugated secondary antibodies (goat anti-mouse, Life Technologies) and 5 μg/ml Alexa 647-conjugated SNA. The cells were washed 3 times in PBS and post-fixed in freshly prepared 4 % paraformaldehyde for 10 min at RT. A549 cells in Fig. 1B were labelled with 10 ng/ml SNA-Alexa647 and 1μg/ml Hoechst33342 (Life Technologies) in DMEM.

### SMLM microscopy

EGFR single- and two-color STORM imaging were performed using a recently developed flat-field epi illumination microscope [58]. Briefly, two lasers with wavelengths of 642 nm (2RU-VFL-P-2000-642-B1R, MPB Communications) and 750 nm (2RU-VFL-P-500-750-B1R, MPB Communications) were used to switch off fluorophores on the sample, while a 405 nm laser (OBIS, Coherent) controlled the return rate of the fluorophores to the fluorescence-emitting state. A custom dichroic (ZT405/561/642/750/850rpc, Chroma) reflected the laser light and transmitted fluorescence emission before and after passing through the objective (CFI60 PlanApo Lambda Å~60/NA 1.4, Nikon). After passing the respective filter (ET700/75M, Chroma or ET810/90m, Chroma), emitted light from the sample was imaged onto the sCMOS camera (Zyla 4.2, Andor). Axial sample position was controlled using the pgFocus open hardware autofocus module (http://big.umassmed.edu/wiki/index.php/PgFocus). Typically, 20,000 frames at 10 ms exposure time were recorded using Micromanager[59]. Imaging was performed using an optimized STORM buffer as described previously[60]. Image stacks were analyzed using a custom CMOS-adapted analysis routine[61]. Lateral sample drift was corrected using either image correlation (Thunderstorm[62]) or gold fiducial markers (B-Store, https://github.com/kmdouglass/bstore). Two-color datasets were analyzed using LAMA[63]. Random datasets for CBC analysis were generated in MatLab.

SNA single color imaging was performed on a modified Olympus IX71 inverted microscope. A 641 nm laser (Coherent, CUBE 640-100C) and a 405 nm laser (Coherent, CUBE 405-100C) was reflected by a multiband dichroic (89100 bs, Chroma) on the back aperture of a 100x 1.3 NA oil objective (Olympus, UplanFL) mounted on a piezo objective scanner (P-725 PIFOC, Physik Instrumente). The collected fluorescence was filtered using a band-pass emission filter (ET700/75, Chroma) and imaged onto an EMCCD camera (IxonEM+, Andor) with a 100 nm pixel size and using the conventional CCD amplifier at a frame rate of 25 fps. Laser intensity on the sample measured after the objective was 2 - 4 kW/cm^2^. 20,000 frames at 30 ms exposure time were recorded using Micromanager[59]. Image stacks were analyzed using ThunderStorm[62]. Lateral sample drift was corrected using either image correlation (Thunderstorm[62]) or gold fiducial markers (PeakSelektor, IDL, courtesy of Harald Hess).

PALM imaging was performed on a Zeiss Axio Observer D1 inverted microscope, equipped with a 100x, 1.49 NA objective (Zeiss). Activation and excitation lasers with wavelengths 405 nm (Coherent cube) and 561 nm (Crystal laser) illuminated the sample in total internal fluorescence (TIRF) mode. We used a four color dichroic 89100bs (Chroma), fluorescence emission was filtered with an emission filter ET605/70 (Chroma) and detected with an electron-multiplying CCD camera (iXon+, Andor Technology) with a resulting pixel size of 160nm. For each region of interest, typically 10000 images of a 41×41 μm2 area were collected with an exposure time of 30 ms. Photoactivatable proteins were activated with 405 nm laser intensity < 0.5 W/cm^2^, chosen to maintain a sparse population of activated molecules for localization, and excited with 561 nm laser intensity of ∼1 kW/cm^2^. Image stacks were analyzed using ThunderStorm[62].

### STED microscopy

STED measurements were done with Abberior STED microscope (Abberior Instruments, Germany) as previously described in [64,65]. The microscope is equipped with a titanium-sapphire STED laser (MaiTai HP, Spectra-Newport). The labelled Abberior Star Red-labelled SNA was excited using 640 nm pulsed diode laser (Picoquant, Germany) with an average excitation power of 5-10 μW at the objective (UPlanSApo 100x/1.4 oil, Olympus). Depletion was achieved using tunable pulsed laser at 780 nm. The microscope was operated using Abberior’s Imspector software.

### Cluster analysis

For the cluster analysis, we used the algorithm density-based spatial clustering applications with noise (DBSCAN)[66], which was embedded into our custom analysis MatLab routine. DBSCAN only needs two input parameters, *Eps* and *k*. It then counts for each localization, how many neighbor localization are within a circle of radius *Eps*. If the localization has *k* neighbors within *Eps*, it is classified as part of a cluster. If it does not have enough neighbors within *Eps*, but is itself a neighbor of a cluster localization, it is classified as an edge point. All remaining localization are classified as unclustered. In order to analyze the very dense and heterogeneous localization maps we obtained from SNA imaging, we performed two consecutive DBSCAN runs with different parameters for *Eps* and *k*. Only this allowed us to account for all visually visible clusters. Clustered and edge points are then combined and handed over to the single cluster analysis part of the analysis routine. For each cluster, we examined a set of parameters such as area and mean diameter as well as the number of localizations per cluster. We further analyzed the localization density distribution per cluster by performing a nearest-neighbor search using a search radius of 20 nm. All localization processing was performed using custom written MatLab (MathWorks) scripts.

### Single molecule calibration

In order to measure the localization precision of our system and calibrate the grouping parameters, we performed STORM imaging on isolated dye molecules. 25 mm round glass slides (Menzel, # 1.5) were plasma cleaned for 10 min and coated with poly-L-lysine solution (100 μg/ml in ddH_2_O) for 1 h at room temperature. After washing in ddH_2_O, the slides were dried and incubated with 10 - 50 pM dye-conjugated SNA or anti-EGFR antibodies respectively. After 15 min, the slides were washed once and then imaged under experimental conditions. Localization maps were filtered and drift-corrected using gold fiducials. Individual localizations were first grouped with *gap* = 0 and search radius 30 nm, to merge individual blink events, then grouped again with a gap time equal to the total acquisition time (15k frames). This allows to quantify the spread of localizations along x and y (i.e. the localization precision σ) as well as the time between individual blink events (i.e. the dark time). All localization processing was performed using custom written MatLab (MathWorks) scripts.

### Virus binding simulations

We simulated a flat patch of cell surface (1×1 μm^2^) including *n* attachment factor molecules at random positions. For the analysis of the degree of clustering (Fig. S10B), the simulated molecules were gradually shifted into clusters, while *n* was kept constant. To analyze the impact of the cluster size (i.e. the number of receptors per cluster), we simulated a constant concentration of attachment factor molecules and added receptor clusters at the indicated size (Fig. S10A). A virus attempting to attach was simulated as s two-dimensional projection of a spherical 100 nm virus particle. The virus center was randomly placed onto the simulated surface and the number of attachment factor molecules within a radius of 50 nm was counted. More than 10 molecules were counted as successful binding. Matlab scripts to run the simulation are available at GitHub (https://github.com/christian-7/Virus_Binding_Simulation).

## Acknowledgements

We would like to thank Katharina Horst for support and Andreas Herrmann for helpful discussions on the manuscript.

## Supporting Information

### Supplementary note 1

#### Establishment of the experimental system

It was shown previously that influenza A virus (IAV) activates and uses EGFR to trigger endocytosis and enter into mammalian host cells [9]. Specifically, this was shown for different virus strains including H1N1/PR8, H7N1/FPV and H3N2/X31. To test if IAV H3N2/X31 in our hands entered cells in an EGFR-mediated way, we stimulated human A549 cells with 100 ng/ml EGF, causing EGFR internalization and thereby removal from the cell surface [9]. We found that successful virus infection, as detected by viral nucleoprotein production, was decreased by 40 % as compared to the control. As expected, the effect of completely removing the primary AF sialic acid using sialidase (neuraminidase, NA) treatment was much stronger and reduced the amount of infected cells by 80 % (Fig. S1).

**S1 Figure: IAV infection efficiency in A549 cells is reduced after EGF stimulation or NA treatment.** A549 cells were either treated with 100 ng/ml EGF for 30 min to reduce the concentration of available EGF receptors or 0.01U/ml neuraminidase for 3h at 37 °C. Cells were infected with influenza A/X31 (MOI ∼ 1) for 5h then fixed and immunolabelled for newly produced viral nucleoprotein (NP). The cell nuclei were counterstained with DAPI. Nuclear NP signal was quantified using automated image analysis with Cellprofiler [55].

**S2 Figure: Ezrin labelling and microvilli identification in STORM localization maps**. (**A**) A549 cells were immunolabelled for the actin-binding protein Ezrin, which was shown to be enriched in microvilli [19]. The cells were imaged using STORM. Microvilli are clearly distinguishable and resemble the large cluster population observed in SNA labelled cells as well as observations from scanning electron microscopy (SEM, inset). Scale bars: left panel: 2 μm, right panel: 500 nm, inset: 200nm. (**B**) Ezrin localization maps can be used to set a threshold for the clusters size obtained from DBSCAN clustering to specifically analyze the not-microvilli cluster population in SNA localization maps (Fig. 2).

**S3 Figure: Experimentally obtained localization precision *σ* for Alexa 647 and Alexa 750**. Glass slides were washed, plasma cleaned and coated with Poly-L-lysine (0.01 % in water) for 1h. Conjugated antibodies were diluted in PBS to a final concentration of ∼10 nM and adsorbed to the coated glass slides. Individual molecules were imaged under experimental conditions. Localizations originating from single Alexa 647 (**A**) and Alexa 750 (**B**) molecules were aligned to allow the estimation of the average localization precision: σ_x,y_A647 = 12 nm and σ_x,y_ A750 = 21 nm.

**S4 Figure: Localization precision partly mimics local concentration gradient.** To test if the localization precision accounts for the gradient in localization density we observed in AF clusters (Fig. 3), we simulated clusters of random localizations (**A**) using cluster size data taken from our experimental STORM measurements (i.e. radius r, number of localizations *n*, localization precision σ). The local density was then determined using σ nearest neighbor search within a radius of 3σ. We indeed observed that the simulated clusters exhibit an up to about 8-fold local enrichment (see one example in **A-C**). We then simulated clusters following the full distribution of experimental data (i.e. radius *r*, number of localizations *n*). Comparing with the density gradient observed in our experimental data (**D** and Fig.3), we find that both distributions are well separated and that the described effect only accounts for density changes < 8-fold.

**S5 Figure: IAV single-virus tracking on A549 cells.** Single virus tracking on live A549 cells revealed four main types of virus movement: (**A**) three-stage movement, (**B**) confined, (**C**) mixed, (**D**) drift. The fraction of all modes of movement was analyzed at the indicated conditions (**E**).

**S6 Figure: Molecule blinking correction for EGFR data.** To estimate the number of emitting molecules from a STORM dataset and to avoid false clustering of individual molecules, we merged multiple localizations originating from the same molecule into a single localization. The merging procedure requires a gap distance as well as a gap time, within which localizations will be counted as originating from the same molecule. To calibrate these values, we imaged isolated labelled anti-EGFR antibodies under experimental conditions. Localizations originating from a single molecule could be grouped to determine their lateral spread (**A**) as well as the dark time between individual bursts (**B**). To ensure a high certainty of merging, the dark time cut-off was determined by the 99 % quantile to 18 s. Using the experimentally determined spread of localization (**A**, 35 nm) and the dark-time cut off, localization bursts from the same molecule can now be combined into a single position. While each molecule is counted multiple times due to molecule blinking (uncorrected, **C**), merging allows a more precise estimate of the molecule numbers while avoiding false clustering (corrected, **D**).

**S7 Figure: Experimental positive control for CBC-based colocalization analysis.** We used two differently labelled versions of SNA as an experimental colocalization positive control. This served us also as a nominator to better evaluate the degree of colocalization of our test molecule pair SNA/EGFR. A549 cells were labelled with two SNA variants, conjugated to Alexa 647 as well Alexa 555 (**A**). Both localization datasets were analyzed using CBC resulting in a colocalization value C_A_ associated to each individual molecule. A histogram of C_A_ for one channel is shown in **B**. We set the threshold to 0.3, above which localizations were counted as colocalized. **C** shows one SNA dataset color coded according to C_A_. **D** shows all localizations from the same dataset with C_A_ > 0.3.

**S8 Figure: EGFR cluster size and molecule number after IAV and EGF stimulation.** A549 were either left in culture medium (control) or stimulated with IAV or 100 ng/ml EGF for 15 min. The cells were fixed and immunostained using anti-EGFR antibodies. Upon either stimulation, we could not detect a change in the size of the EGFR clusters (**A**) or the amount of molecules per cluster (**B**). **B**, legend as in **A**.

**S9 Figure: Single molecule diffusion coefficients from sptPALM of EGFR-mEos3 in A549 cells.** A549 cells were transiently transfected with EGFR-mEos3. sptPALM imaging and analysis of single molecule trajectories revealed a wide range of diffusion coefficients. The left panels in **A** and **B** show the distribution of diffusion coefficients from sptPALM obtained at the apical (**A**) and the basolateral plasma membrane (**B**). Molecules were classified as mobile (D>0.5 μm2/s) or immobile (D<0.5 μm2/s) respectively. Calculated MDS plots (**A** and **B**, right panels) for both classes exhibit a rather linear dependence for the mobile fraction, while the curve saturates with increasing lag time for the immobile fraction, the latter indicating spatial confinement.

**S10 Figure: Virus Binding Simulation.** To estimate the effect of AF clustering on the efficiency of a virus to bind the target cell, we simulated two scenarios in a 1×1 μm membrane area, (**A**) a varying cluster size and (**B**) a varying degree of clustering. For **A**, we simulated a constant lateral concentration of AF (black) and added AF clusters (blue) at increasing size. In **B**, we keep the total amount of AF constant and gradually shift molecules into clusters. In both cases, an approaching virus was simulated as a 2D projection of a small spherical IAV particle (contact area as red circles in **A** and **B**). A binding attempt was counted as successful if at least 10 AF molecules were found inside the contact area. **C** and **D** show the simulation result plotted as the binding probability out of 1000 simulations against the respective tested cluster parameter.

**S11 Figure: EGFR activation upon SNA binding and pharmacological EGFR cluster disruption in A549 cells.** (**A**) We wanted to test if SNA (tetramer), which has a lower binding valency against sialic acid compared to IAV, is still able to activate EGFR in A549 cells. To this end, cells were either stimulated with EGF or incubated with DMEM supplemented with 50 μg/ml unlabelled SNA. To diminish the SNA-SA interaction, we also treated the cells with 0.1 units/ml sialidase for 90 min before SNA stimulation. The cells were fixed and immunolabelled using antibodies against phopho-EGFR (Y1068). Although, we detected an increase in the phopho-EGFR cluster density upon EGF stimulation, we did not detect a difference after incubation with SNA. (**B**) The stability of EGFR clusters was tested upon pharmacological cell treatment either to inhibit actin polymerization using latrunculin A (0.2 μM for 30 min) or cholesterol depletion using methyl-β-cyclodextrin (40 μg for 60 min). After the treatment, the cells were fixed and immunolabelled using antibodies against EGFR. Following both treatment, we detected a decrease in the clustered fraction suggesting cluster destabilization.

**S1 Video: Evolution of the confinement index I_conf_ as well as the particles distance from the origin for a simulated virus trajectory.** The particle moves according to D_free_ until it encounters an AF cluster (top panel, red circles). Due to the higher concentration of AF, the particles diffusion is slowed down to D_conf_ and the particle is confined. Particle confinement is detected by an increase of the confinement index (middle panel) as well as a plateau in the distance from origin plot (lower panel).

